# Millisecond nonlinear state changes during droplet coalescence identify therapeutic-antibody developability liabilities

**DOI:** 10.64898/2026.05.06.723251

**Authors:** Alexander Nicholas St John, Jack Holland, Elizabeth Suk-Hang Lam, Sejeong Lee, Piergiorgio Caramazza, Alec Nelson Thomas, Shamit Shrivastava

## Abstract

Apoha’s Liquid State Intelligence Platform (LSIP) records ellipsometric waveforms from injections depositing sub-microgram quantities of antibody drop-by-drop onto a liquid reservoir. We previously showed that a behavioural feature extracted from the waveforms, VIBE1, identified antibodies carrying multiple biophysical liabilities in an industrial dataset of 71 monoclonal antibodies, and enriched for clinical failure across a larger dataset of 235 therapeutic antibodies [1]. Here, we use an auxiliary coalescence-sensor channel to decode VIBE1 by separating the coalescence event from its propagation through the substrate. The per-titration drop-to-drop standard deviation of pinch-off time, *σ*_*τ*_, explains most of VIBE1’s variance across the dataset (*R*^2^ = 0.92, *n* = 1182). High-speed imaging at 10,000 frames per second reveals that all imaged drops initially thin at the same Newtonian capillary–inertial rate while the neck remains wide. In drops from certain antibodies, the thinning bridge then decelerates as internal strain builds in the narrowing neck. This elasto-capillary stiffening response has a timescale *λ* that decreases as pinch-off time *τ*_*i*_ increases across the imaged set. *σ*_*τ*_ is therefore a readout of the antibody’s propensity to undergo a transient gel-like stiffening response during coalescence, and that variability is what VIBE1 captures. The signal is concentration dependent, and absent in bovine serum albumin (BSA) tested at up to an order of magnitude higher molarity than the antibodies, despite BSA being a strongly surface-active globular protein. The instrument is configured so that complex behaviours of this kind appear in its recorded waveforms; the gel-like coalescence response we identify here is one such phenomenon.

---

The behaviour of therapeutic antibodies at interfaces and under mechanical stress is a recognised driver of aggregation, particle formation, and accelerated clearance [2, 3, 4]. Antibodies adsorbed to the air–water interface form glassy, soft-solid films at concentrations relevant to manufacturing and dispensing [4, 5], and adsorption to hydrophobic surfaces produces irreversible globular layers [6]. Yet the stresses encountered by a therapeutic in its life cycle are not the single-axis perturbations isolated by conventional assays: real failure environments compose adsorption, compression, extensional flow, and chemical-environment shifts on millisecond-to-second timescales [3, 5, 7, 8], acting synergistically on molecular structure. The properties through which they act are themselves context-dependent: hydrophobicity is the design principle of pulmonary surfactant proteins [9], conformational lability initiates haemostasis through von Willebrand factor [10], and antibody CDRs themselves rely on hydrophobic and conformationally adaptive residues to bind antigen [11, 12]. The canonical early-stage developability panels (hydrophobic interaction chromatography, self-interaction nanoparticle spectroscopy, differential scanning calorimetry, polyreactivity assays) each capture a single biophysical axis [13, 14, 15] and do not by themselves identify which combinations of property values constitute a liability under composite stress. Direct integrated stress assays — agitation, shake, foam, freeze–thaw — probe the composite response, but typically require 100–1000 mg of material per condition over multi-day protocols and on formulated drug product [2, 16], deferring this assessment to the stage of development at which the cost of correcting a liability is already high.

We recently introduced Apoha’s Liquid State Intelligence Platform (LSIP) and the VIBE (Variations in Interfacial Behaviour upon Excitation) family of behavioural features, which address both problems together [1]. Antibodies are delivered drop-by-drop into a near-thermodynamic-transition liquid substrate at sub-10 µg per measurement, and the resulting interfacial waves are recorded ellipsometrically across ∼100 successive coalescence events per titration. The primary feature, VIBE1, identified antibodies carrying multiple biophysical liabilities in an industrial dataset of 71 molecules and enriched for clinical failure across a 235-mAb clinical-stage dataset, while correlating only modestly with any individual conventional assay, indicating a dependence on a combination of multiple properties rather than any single property in isolation. VIBE1 is therefore empirically useful but mechanistically unexplained. A physical account matters for two reasons: interpretability strengthens the empirical result by placing VIBE1 on the same footing as the conventional axes it integrates over, and the principle of reading a molecule’s integrated stress response from its signature at a coalescing interface should generalise beyond antibodies if its physical basis can be articulated.

Here we use the instrument’s auxiliary coalescence sensor to decode what VIBE1 measures. The combination of attributes VIBE1 responds to turns out to be a stochastic, antibody-specific transition to a transient gel-like phase during droplet coalescence, observed directly by high-speed imaging and quantitatively linked to the dataset-scale signal. By separating the coalescence event itself from its propagation through the substrate, we find that a single per-titration feature of the coalescence channel, the drop-to-drop standard deviation of pinch-off time *σ*_*τ*_, explains *R*^2^ = 0.92 of VIBE1 across our 1182-titration dataset [1], and correlates more strongly with most of the conventional developability panel than VIBE1. To ground that population observable in a physical event, we image individual coalescence events at 10,000 frames per second (fps). All drops, water and antibodies, share the same Newtonian capillary–inertial rate when the bridge is wide; in some drops the bridge then decelerates as internal strain rises in the narrowing neck, with an elasto-capillary stiffening timescale *λ* that tracks *τ*_*i*_ inversely across the imaged set. *σ*_*τ*_ can therefore be interpreted as the antibody’s propensity to undergo this gel-like transition, and VIBE1 encodes this propensity.

## Results

### Drop-to-drop pinch-off variability is the dominant proximate cause of VIBE1

Apoha’s LSIP reads out a per-droplet response from a small analyte injection deposited drop-by-drop into a sensing-liquid reservoir. Each deposition is recorded on multiple physically distinct channels (Figure 1). The instrument’s primary readout is a set of ellipsometric channels measuring the response of the sensing-liquid surface, an aqueous reservoir containing an anionic surfactant held near its critical micellisation concentration, to the coalescence event at a fixed distance from the coalescence point. An auxiliary upstream coalescence sensor (a laser beam below the dispenser, partially occluded by the dispensing droplet) tracks the droplet itself as it approaches, contacts, and separates from the substrate. We focus here on one of the ellipsometric channels, *S*_Y_, which encodes both spatial and temporal information about the coalescence event and information about the substrate the wave then propagates through.

**Figure 1:**
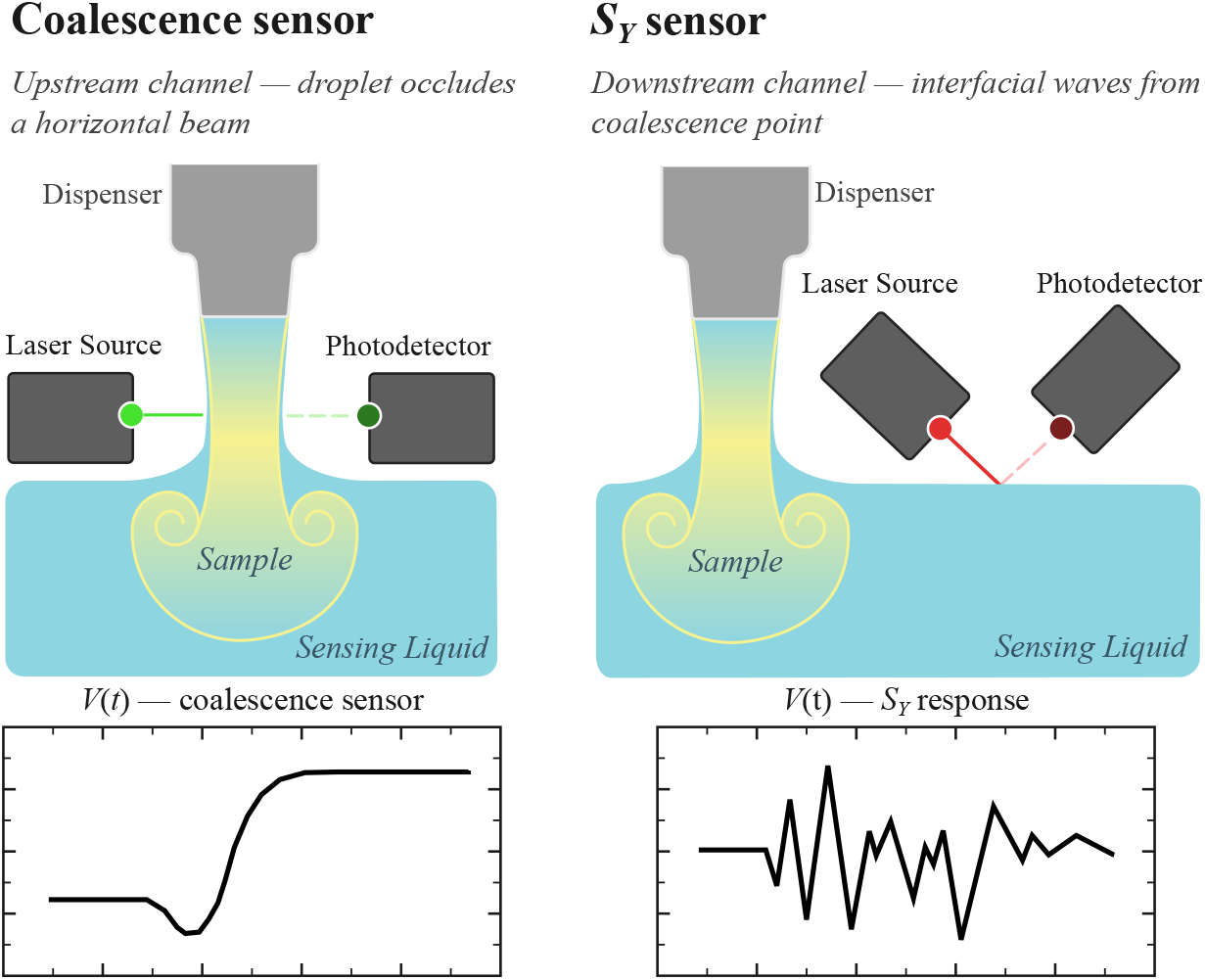
Measurement architecture of the LSIP. The instrument records multiple physically distinct channels per droplet deposition; the two channels used in this paper are shown. (Left) The auxiliary coalescence sensor: a horizontal laser beam below the dispenser, partially occluded by the dispensing droplet, records a waveform *V* (*t*) that tracks the droplet itself as it approaches, contacts, and separates from the sensing-liquid surface. (Right) The primary readout: an ellipsometric channel *S*_Y_ that reports the response of the sensing-liquid surface to the coalescence event, recorded at a fixed distance from the coalescence point. VIBE1 is computed from *S*_Y_. The two channels sit in a defined causal order: the upstream sensor records the coalescence event at its source; *S*_Y_ records its downstream consequence as the resulting mechanical perturbation propagates across the substrate.

In our previous work, we took a top-down approach to identifying behavioural features from the response channels [1]. A set of waveform statistics (the VIBEs) were extracted to capture discrete spatio-temporal distributions of waveforms, screened for discriminatory power across the panel of 235 clinical-stage monoclonal antibodies, and identified VIBE1 as the most discriminatory single feature. VIBE1 is the per-titration mean kurtosis of the *S*_Y_ waveform within the 104–140 ms window of each drop, evaluated across drops 20–40 of the titration:

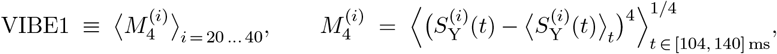

where the inner average is over time within the window. Antibodies with high VIBE1 scores carry significantly more biophysical liabilities across a panel of conventional assays. The feature is therefore empirically useful but not mechanistically explained: it correlates only weakly and simultaneously with hydrophobicity, polyreactivity, and thermal-stability assays, with no single conventional axis dominating.

To begin to identify the physical content of VIBE1 we ask how much of its variance is set by the coalescence event itself, as distinct from the propagation response of the substrate. The two channels of Figure 1 sit in a defined causal order: the upstream sensor records the coalescence at its source; *S*_Y_ records its downstream consequence as the resulting wave propagates across the substrate. Any antibody-dependent feature of *S*_Y_ therefore originates in the coalescence event itself, in the propagation medium, or in both.

Successive drops within a single titration produce nearly identical coalescence-sensor traces, except for the timing of a secondary peak in the time-derivative of the signal, which jitters between drops (Figure 2A,B). This feature corresponds to rupture of the liquid bridge connecting the dispensing tip to the coalesced droplet: as the bridge thins, surface tension drives neck collapse via the Rayleigh–Plateau instability [17, 18] on a capillary–inertial timescale of 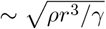, and the rupture itself is the most abrupt mechanical event of the deposition that results in a decrease of light absorbed. We identify this secondary gradient maximum as the pinch-off time

**Figure 2:**
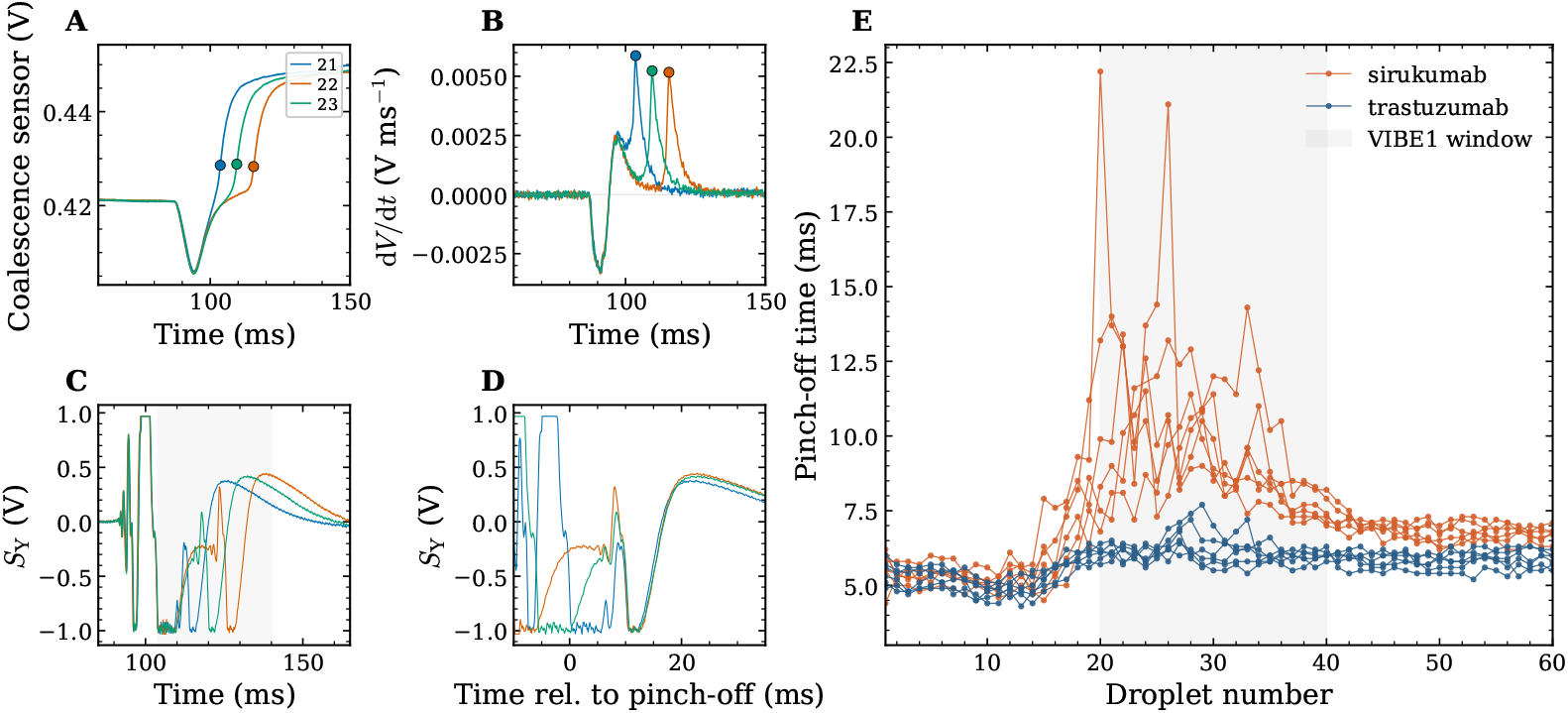
Pinch-off variability causally drives VIBE1. (A) Coalescence-sensor traces for three successive drops from a single high-VIBE1 titration; the traces overlap except for the timing of bridge rupture (markers, *τ*_*i*_). (B) Smoothed time-derivative of the same traces; the secondary gradient maximum identifies *τ*_*i*_, with per-titration drop-to-drop standard deviation *σ*_*τ*_. (C) Corresponding *S*_Y_ waveforms (the primary, ellipsometric readout) in absolute time. A second pulse appears beyond the common initial transient (*t<* 110 ms; VIBE1 window 104–140 ms shaded), with arrival time varying in lockstep with *τ*_*i*_. (D) Aligning the same waveforms to *τ*_*i*_ collapses this second pulse onto a common shape (∼4 ms fixed propagation delay): the late-arriving *S*_Y_ pulse is a stereotyped consequence of pinch-off. (E) Per-drop *τ*_*i*_ across the deposition for six titrations of sirukumab (orange, high VIBE1) and six of trastuzumab (blue, low VIBE1); shaded VIBE1 analysis window (drops 20–40). The VIBE1-window exclusion (Methods) removes 22 of ∼24,800 drops across the dataset and does not affect the data shown here. Sirukumab spans ∼5–20 ms within the window, with consecutive drops jumping by several-fold (not Gaussian noise around a smooth response but flipping between regimes), while trastuzumab is tightly clustered near baseline. Across the dataset, *σ*_*τ*_ explains *R*^2^ = 0.92 of VIBE1 (Pearson, *n* = 1182; Figure 3A).

*τ*_*i*_ for the *i*th drop, and define

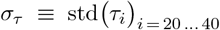

across drops 20–40 (the VIBE1 analysis window) as a per-titration feature of the coalescence event itself, independent of the substrate’s response. High-speed imaging confirms the identification (next section).

The *S*_Y_ waveforms have two regimes (Figure 2C). Before ∼110 ms, all drops in a titration produce nearly identical waveforms, the common initial coalescence transient. Beyond ∼110 ms, a second oscillatory pulse appears whose arrival time varies between drops, matching with a fixed offset the variability in *τ*_*i*_. Aligning each *S*_Y_ trace on its *τ*_*i*_ collapses this second pulse onto a common shape (Figure 2D), with a ∼4 ms residual offset consistent with a constant propagation delay from the pinch-off site to the sensor. The late *S*_Y_ pulse, the part of the waveform from which VIBE1 is computed, is dominated by *τ*_*i*_. Per-drop *τ*_*i*_ itself varies widely at fixed bulk concentration: across six sirukumab titrations (high VIBE1) it spans ∼5–20 ms within the analysis window, with consecutive drops jumping by several-fold (Figure 2E); across six trastuzumab titrations (low VIBE1) it stays tightly clustered near baseline. High-speed imaging (next section) shows the variability comes from the bridge-thinning kinematics themselves.

After excluding drops whose late *S*_Y_ pulse falls outside the 104–140 ms window (Methods; 22 of ∼24,800, 0.09%), *σ*_*τ*_ explains *R*^2^ = 0.92 of VIBE1 across the 1182-titration dataset (Pearson, *n* = 1182; Figure 3A): drop-to-drop pinch-off variability accounts for 92% of VIBE1’s variance across the dataset. The relationship is preserved when within-antibody replicate variability is averaged out: per-antibody mean *σ*_*τ*_ versus per-antibody mean VIBE1 gives *R*^2^ = 0.96 across *n* = 289 antibodies (Supplementary Fig. S5). The 235 antibodies were also characterised against the 11-assay Jain panel [13, 14] (thermal stability *T*_*m*_, accelerated-stress aggregation AS, salt-gradient affinity capture SGAC, hydrophobic interaction chromatography HIC, self-interaction-spectroscopy variants SMAC, ACSINS, and CSI, polyspecificity reagent binding PSR, cross-interaction chromatography CIC, non-specific ELISA binding, and baculovirus-particle binding BVP). Per-antibody Spearman *ρ* against each assay (Figure 3B) is larger in absolute value for *σ*_*τ*_ than for VIBE1 on most axes. Therefore, the physical observable derived by decoding VIBE1 correlates more strongly with the conventional developability panel than the statistical summary it was decoded from.

**Figure 3:**
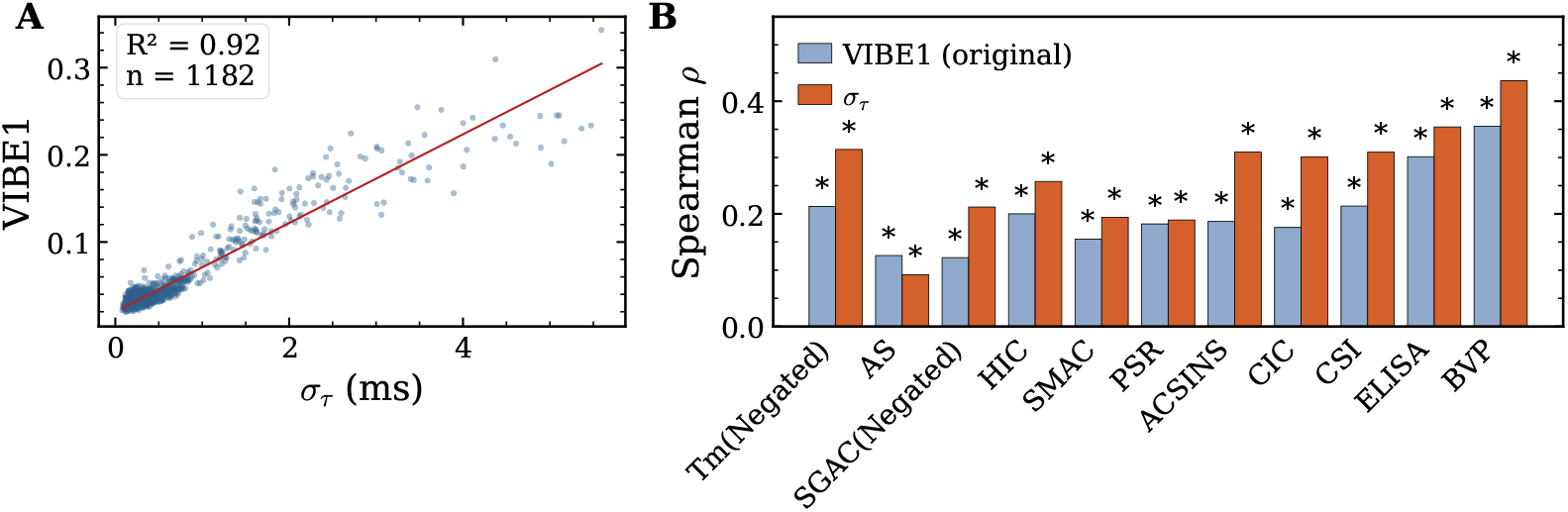
*σ*_*τ*_ explains VIBE1 across the dataset and correlates more strongly than VIBE1 with the conventional developability panel. (A) Per-titration *σ*_*τ*_ vs. per-titration VIBE1 (1182 titrations, VIBE1-window exclusion applied; Methods). Pearson *R*^2^ = 0.92, *n* = 1182. (B) Per-antibody Spearman *ρ* between the 11 biophysical assays and either VIBE1 (blue) or *σ*_*τ*_ (orange, per-antibody mean). Asterisks mark *p <* 0.05. Most assays show a larger |*ρ*| for *σ*_*τ*_ than for VIBE1. Assay abbreviations: *T*_*m*_, thermal stability; AS, accelerated-stress aggregation; SGAC, salt-gradient affinity capture; HIC, hydrophobic interaction chromatography; SMAC, ACSINS, CSI, self-interaction-spectroscopy variants; PSR, polyspecificity reagent binding; CIC, cross-interaction chromatography; ELISA, non-specific antigen binding; BVP, baculovirus-particle binding.

### High-speed imaging directly links pinch-off to VIBE1 and resolves the underlying elasto-capillary stiffening response

To link bridge pinch-off to VIBE1 directly, we imaged single coalescence events at 10,000 fps on the same instrument: 12 drops in total (1 water, 2 trastuzumab, 2 galiximab, 7 fresolimumab) chosen to span the dataset’s VIBE1 range (trastuzumab, low VIBE1; fresolimumab and galiximab, high VIBE1). The choice reflects each antibody’s VIBE1 ranking in the dataset, not a per-molecule clinical-failure attribution. The camera is used here as a per-event confirmatory probe; dataset-level statistics are in the previous section and in ref. [1]. As shown below, all drops thin at the same Newtonian capillary–inertial rate when the bridge is wide; in the high-VIBE1 antibodies the bridge then enters an exponential, viscoelastic-like thinning regime as internal strain rises in the narrowing neck, and pinches off later than the water and low-VIBE1 controls.

The water-only video (Figure 4) resolves five stages: approach before contact; a sharp drop in transmitted intensity at droplet–substrate contact; a brief expansion phase in which the bridge between dispensing tip and substrate grows to its maximum radius; continuous thinning of the bridge as surface tension drives the neck towards collapse; and rupture of the bridge separating the dispensing tip from the deposited droplet (Figure 4A). The frame-by-frame mean image intensity tracks the upstream-detector signal closely, and the time-derivative of either trace has the same two-peak structure, with the secondary maximum coinciding with the imaged rupture frame to within ∼100 µs (one frame; Figure 4B). Direct imaging therefore validates the identification of *τ*_*i*_ as bridge-rupture time.

**Figure 4:**
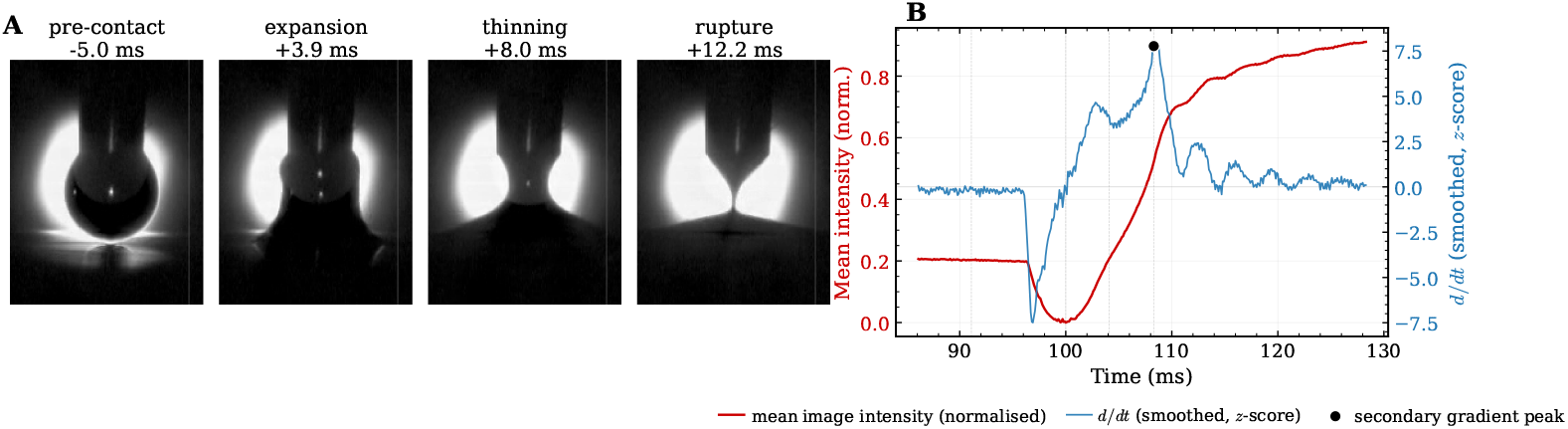
Direct imaging confirms the bridge-rupture event. (A) High-speed video frames from the buffer-only (water) coalescence event, captured at 10,000 fps. Four representative frames are shown, covering the deposition: pre-contact approach, post-contact expansion, mid-thinning of the dispensing-tip liquid bridge, and the rupture frame itself. Frame numbers and acquisition times are indicated; the rupture frame is marked. (B) Mean image intensity (red, normalised to within-event [0, 1]) computed frame-by-frame from the same event, with its smoothed time-derivative overlaid (blue, *z*-scored against the within-event mean and standard deviation). Vertical lines link each panel-A frame to its position on the intensity trace. The black point marks the maximum of the secondary gradient peak, which coincides with the imaged frame of bridge rupture (within one video frame, ∼100 µs).

All twelve drops, water and antibodies alike, share the same early thinning kinematics: when the neck is wide, 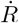 is set by Newtonian capillary–inertial physics and the water and antibody curves overlap within the imaging precision (Figure 6A, early portion of every *R*(*t*) trace). The drops diverge as the neck thins. In some, thinning continues approximately linearly through to rupture; in others the bridge decelerates as internal strain rises in the narrowing neck, following the elasto-capillary form

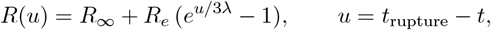

where *u* is the time-to-rupture coordinate. This is the inertio-capillary to elasto-capillary crossover characteristic of low-viscosity elastic fluids [19]. The fitted *λ* is the deceleration timescale: smaller *λ* means stronger late-phase stiffening. The illustrative fresolimumab drop in Figure 5 has *λ* ≈ 2 ms over ∼22 ms of pre-rupture thinning; the illustrative trastuzumab drop reaches rupture in ∼10 ms with no visible deceleration and a correspondingly larger *λ* ≈ 10 ms. Across the 12 drops, *λ* ranges ∼2–10 ms and tracks *τ*_*i*_ inversely (Figure 6B): long-pinch fresolimumab drops have small *λ*; short-pinch water, trastuzumab, and galiximab drops have large *λ*.

**Figure 5:**
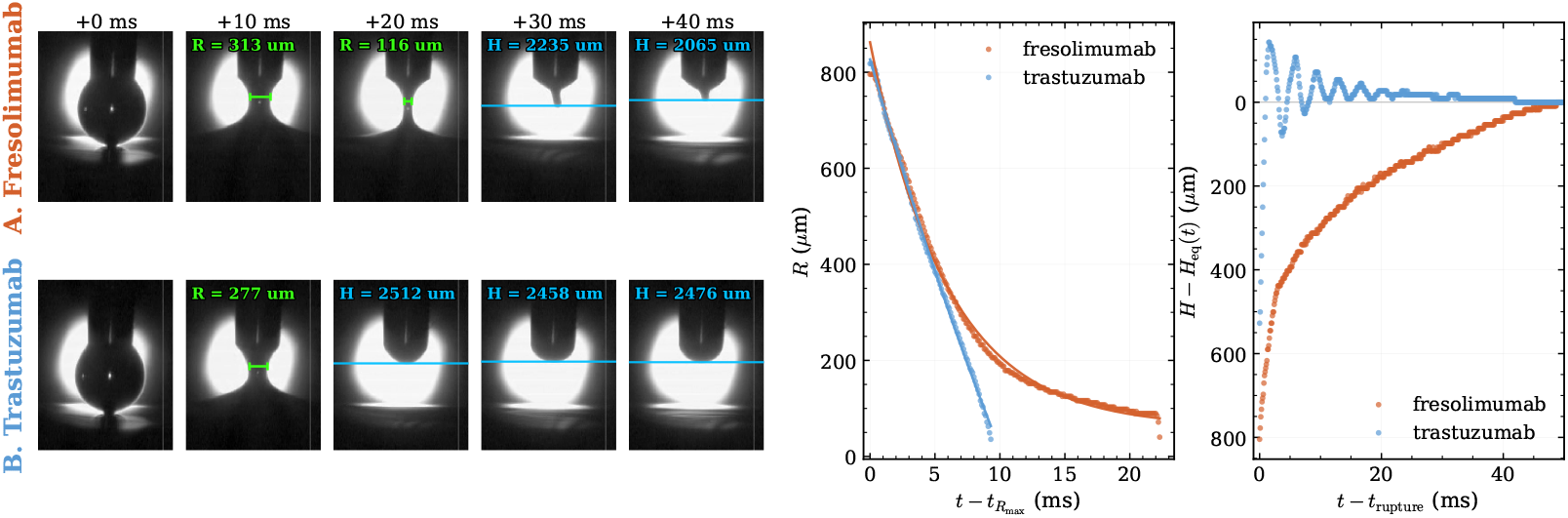
Two illustrative coalescence events: a long-pinch fresolimumab drop with strong late-stage stiffening, and a short-pinch trastuzumab drop without. Both drops were measured at the same 0.1 mg mL^*−*1^ working concentration. (A,B) High-speed video frames at *t* = 0, +10, +20, +30, and +40 ms relative to droplet–substrate contact. (A) Fresolimumab: ∼22 ms of pre-rupture thinning visibly decelerates as the neck narrows; the merged drop persists at a non-spherical shape through the imaging window. (B) Trastuzumab: ∼10 ms of approximately linear thinning followed by free oscillation about the equilibrium spherical shape. Green brackets mark the sub-pixel-refined minimum neck radius *R* during pre-rupture thinning; blue lines mark the bottom-edge height *H* of the merged drop post-rupture. (Right) Pre-rupture neck radius *R*(*t*) (centre) and post-rupture detrended height *H* − *H*_eq_(*t*) (right) for the two drops. Each drop is anchored to its own value of *H* at *t*_rupture_ + 50 ms, so every detrended trace passes through zero at the anchor by construction; oscillatory drops have settled there while non-oscillatory drops appear as a leading deflection that has not yet decayed (Methods). Solid points: smoothed-mask data; solid line on *R*(*t*): per-drop elasto-capillary fit *R*(*u*) = *R*_*∞*_ + *R*_*e*_ (*e*^*u/*3*λ*^ − 1) with *u* = *t*_rupture_ − *t*. The *y*-axis on the *H*(*t*) panel is inverted so that retraction (the upper drop sitting higher than equilibrium) appears as positive deflection. The full videos for the two drops shown are provided as Supplementary Video 6 (fresolimumab) and Supplementary Video 2 (trastuzumab).

**Figure 6:**
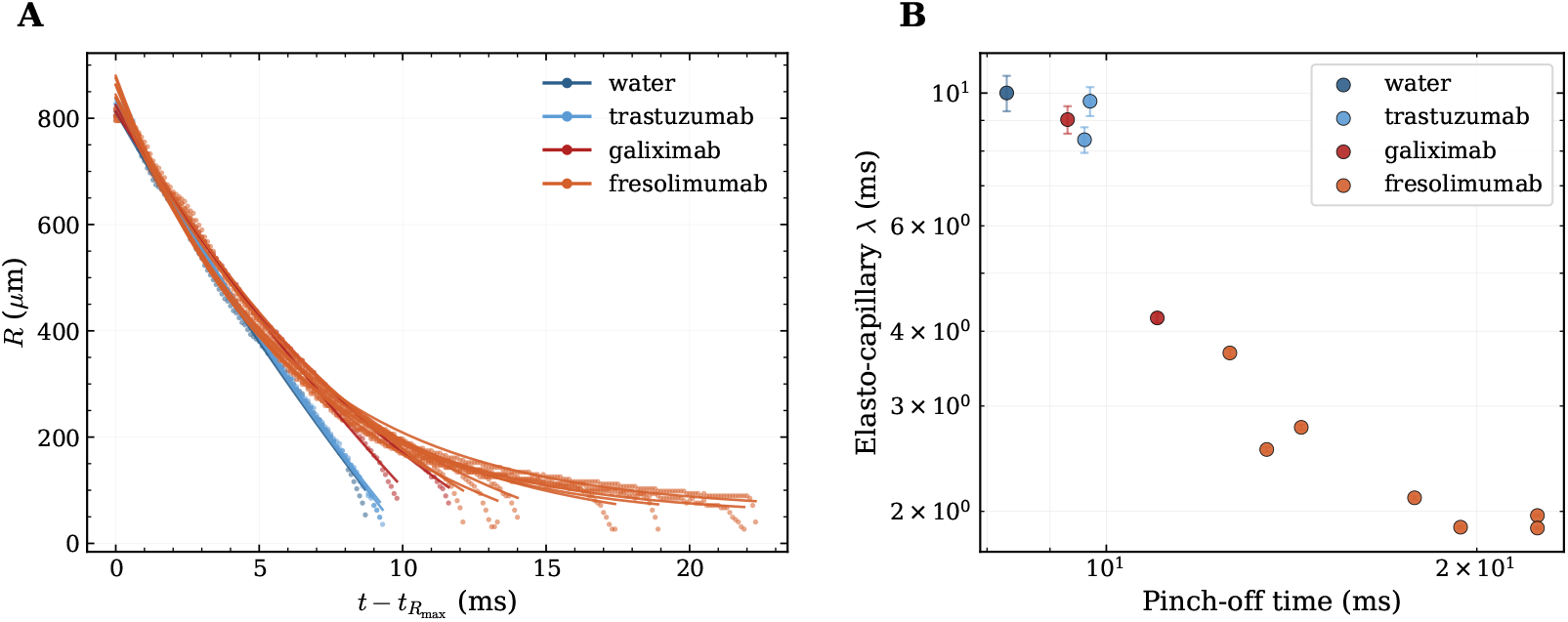
Across three antibodies and a control, longer pinch-off time tracks a smaller elasto-capillary *λ*. (A) Pre-rupture neck radius *R*(*t*) for all 12 imaged drops, anchored at *t* = *R*_max_. Solid points: smoothed-mask data; solid lines: per-drop elasto-capillary fit *R*(*u*) = *R*_*∞*_ + *R*_*e*_(*e*^*u/*3*λ*^ − 1) with *u* = *t*_rupture_ − *t*. Drops coloured by sample (water, trastuzumab, galiximab, fresolimumab). The early portion of every *R*(*t*) curve overlaps within imaging precision; the curves diverge as the neck narrows. (B) Per-drop elasto-capillary *λ* versus pinch-off time *τ*_*i*_ (log-log axes). Vertical bars are sub-pixel-propagated 1 *σ* on *λ. λ* tracks *τ*_*i*_ inversely: long-pinch fresolimumab drops have *λ* ≈ 2 − 4 ms, short-pinch water/trastuzumab drops have *λ* ≈ 8 −10 ms. Per-sample *R*(*t*) and *H* − *H*_eq_(*t*) traces are shown in Supplementary Fig. S1 and Supplementary Fig. S2 respectively.

The post-rupture shape of the merged drop is not part of VIBE1 and is not captured by *σ*_*τ*_: it depends on where the bridge splits and how much of any stiffened phase ends up in the merged drop versus in the reservoir. We report it as a qualitative clue. In drops with limited late-stage stiffening (water, trastuzumab, the short-pinch galiximab drops), the merged drop oscillates freely about its equilibrium spherical shape and returns to within 20% of its initial deflection in 1–4 ms, the standard inertial response of a low-viscosity drop under surface tension. In some long-pinch fresolimumab drops, the merged drop instead persists at a non-spherical shape through the imaging window (Figure 5A; per-sample *H*(*t*) traces in Supplementary Fig. S2). A viscous drop at any plausible IgG-formulation viscosity would relax in well under 5 ms, so the slow relaxation cannot be explained by a homogeneous viscous fluid.

### Concentration response is antibody-specific and not reproduced by a comparably surface-active control

To probe what sets *σ*_*τ*_ we measured its concentration response. Sirukumab and trastuzumab were measured across 0.04–0.25 mg mL^−1^; bovine serum albumin (BSA) at 0.10, 0.20, and 0.50 mg mL^−1^; PBS as the shared *c* = 0 baseline (Figure 7A). Per-drop *τ*_*i*_ traces at the highest tested concentration of each protein are shown in Figure 7B.

**Figure 7:**
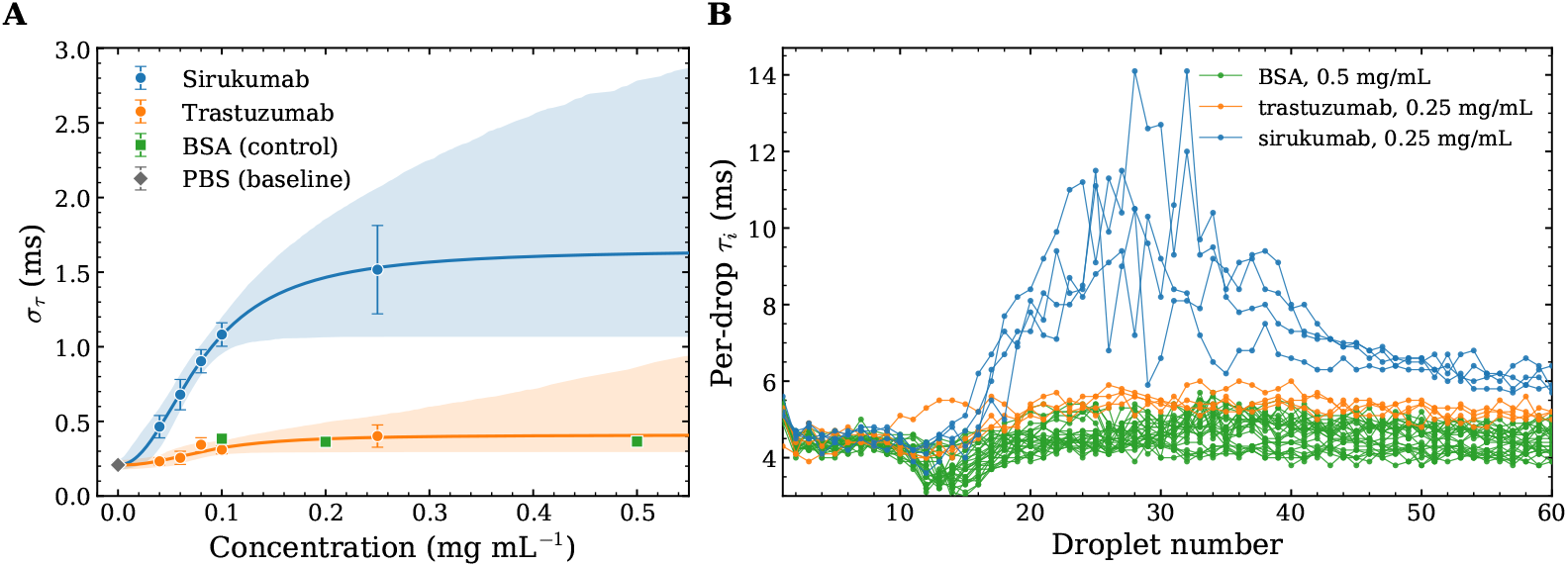
Concentration response: antibody-specific and not reproduced by BSA. (A) Pertitration *σ*_*τ*_ vs. nominal concentration for sirukumab (blue circles), trastuzumab (orange circles), BSA (green squares), and PBS (grey diamond, shared *c* = 0 baseline). At every concentration tested sirukumab is above trastuzumab; BSA tracks just above the PBS baseline across 0.1–0.5 mg mL^*−*1^, 2.3–11-fold above the IgG operating molarity, and below sirukumab at every concentration tested. Solid curves: descriptive Hill curves; shaded bands are 95% confidence intervals from a 2,000-iteration parametric bootstrap. The data are insufficient to constrain *K*_1*/*2_ or Hill *n* quantitatively (Methods). Error bars on points, ± s.d. across replicate titrations. (B) Per-drop *τ*_*i*_ traces vs. droplet number for the highest tested concentration of each protein (sirukumab and trastuzumab at 0.25 mg mL^*−*1^, BSA at 0.5 mg mL^*−*1^); each line is one titration. Sirukumab spans ∼4–14 ms drop-to-drop with consecutive drops jumping by several-fold; trastuzumab stays narrowly distributed near 5 ms; BSA overlaps the trastuzumab traces drop-by-drop. The two antibodies have nominally equal molar mass and are tested under identical conditions, isolating molecule-specific effects from bulk-supply and interfacial-supply artefacts.

Sirukumab and trastuzumab both respond to concentration, but reach very different levels. At 0.10 mg mL^−1^ (∼0.67 µM), sirukumab gives *σ*_*τ*_ ≈ 1.08 ms and trastuzumab gives *σ*_*τ*_ ≈ 0.31 ms, and the gap holds at every concentration tested. Trastuzumab at 0.25 mg mL^−1^, the highest concentration we ran (*σ*_*τ*_ ≈ 0.40 ms), does not reach sirukumab even at 0.06 mg mL^−1^ (*σ*_*τ*_ ≈ 0.68 ms). The per-drop traces in Figure 7B make the same point: sirukumab spans *τ*_*i*_ ∼ 4–14 ms with consecutive drops jumping by several-fold (the regime-flipping pattern from Figure 2E), trastuzumab sits near 5 ms.

BSA does not show a concentration response. From 0.10 to 0.50 mg mL^−1^ (1.5–7.5 µM, 2.3–11× the IgG molarity), *σ*_*τ*_ stays at ∼0.37 ms. The per-drop traces overlap trastuzumab and sit below sirukumab at every concentration we tested.

## Discussion

We set out to explain VIBE1, a per-titration statistic computed from one of the instrument’s ellipsometric channels that enriches for clinical failure across a 235-antibody dataset [1]. By splitting the instrument’s response into the upstream coalescence channel and the downstream ellipsometric channel, we localise VIBE1’s variance to the coalescence event itself rather than to the propagation medium. High-speed imaging at 10,000 fps then shows what is happening at the per-event level, and three features of the imaging together support a transient gel-like phase forming in some drops and not others. First, all twelve imaged drops share the same early thinning kinematics, set by Newtonian capillary–inertial physics; the antibody-specific physics is therefore localised to the late-stage narrowing of the neck rather than to early-stage adsorption or droplet formation. Second, where thinning departs from Newtonian behaviour, the bridge follows an elasto-capillary form with timescale *λ*, and *λ* tracks *τ*_*i*_ inversely across the imaged set: drops that stiffen later in their thinning take longer to rupture. Third, in the strongest cases the merged droplet does not relax to a sphere within the 50 ms imaging window, which a homogeneous viscous fluid at any plausible IgG-formulation viscosity would do in well under 5 ms; an elastic structure with a finite recovery time is the simplest description, consistent with the kinematic signature of arrested coalescence in viscoelastic droplets [20, 21].

The same picture appears at the population level. At fixed bulk concentration, consecutive drops within a single titration switch between long- and short-pinch behaviour (Figure 2E, Figure 7B), pointing to a stochastic, near-bistable transition rather than a graded response. *σ*_*τ*_, the drop-to-drop spread in *τ*_*i*_ at fixed bulk concentration, is therefore the per-event propensity to undergo this transition. Across the 1182-titration dataset, *σ*_*τ*_ explains 92% of VIBE1’s variance and shows higher per-antibody Spearman correlations with the conventional developability panel than VIBE1 itself (Figure 3). *σ*_*τ*_ is a physical observable (drop-to-drop variability of bridge-rupture time) whilst VIBE1 is a statistical summary (kurtosis of *S*_Y_ in a fixed window) of the downstream wave produced by that event. That the physical observable outperforms the statistical summary it was decoded from confirms the decoding that VIBE1 was a proxy for *σ*_*τ*_, and what it reads out is the antibody’s propensity to undergo a transient gel-like transition during coalescence.

Two simple accounts of *σ*_*τ*_ fail against the concentration data. Trastuzumab at our highest concentration does not reach sirukumab at our lowest, ruling out a readout set purely by bulk supply of antibody. BSA forms an interfacial viscoelastic film of its own [22], but at molarities up to tenfold those of the IgGs, gives a flat response, ruling out a readout set by surface activity in the generic sense. The signal therefore reflects something that differs between two IgGs of identical molar mass yet vanishes in a globular monomer. We cannot, from this comparison, separate surface activity per se from the multidomain architecture of an antibody; that distinction will require fragment and non-IgG controls that will be the focus of future work.

A coalescence event on this instrument is not a single-axis perturbation but a composite of three distinct stress classes acting in a defined temporal sequence (Figure 8). During the ∼4 s of droplet growth before contact, molecules adsorb to the expanding air–water interface, where adsorption imposes conformational stress as the protein structure partially unfolds to maximize interfacial contact [4, 5] (i). At rupture and during neck thinning (*t* ≈ 0–20 ms), bulk extensional flow at the thinning neck stretches soluble assemblies [3, 7, 23, 24], the contracting interface compresses the adsorbed layer in-plane [25], and a surface-tension gradient between droplet and reservoir drives Marangoni flow along the neck [23, 26, 27] (ii). Concurrently, surfactant and salt from the sensing-liquid reservoir mix into the molecule-bearing droplet at the contact boundary, shifting the ionic strength, surfactant concentration, and chemical environment the molecule experiences (iii). The same classes of stress are encountered throughout a therapeutic’s life, in manufacturing, formulation, and *in vivo*, and are not isolated by any single-axis assay.

**Figure 8:**
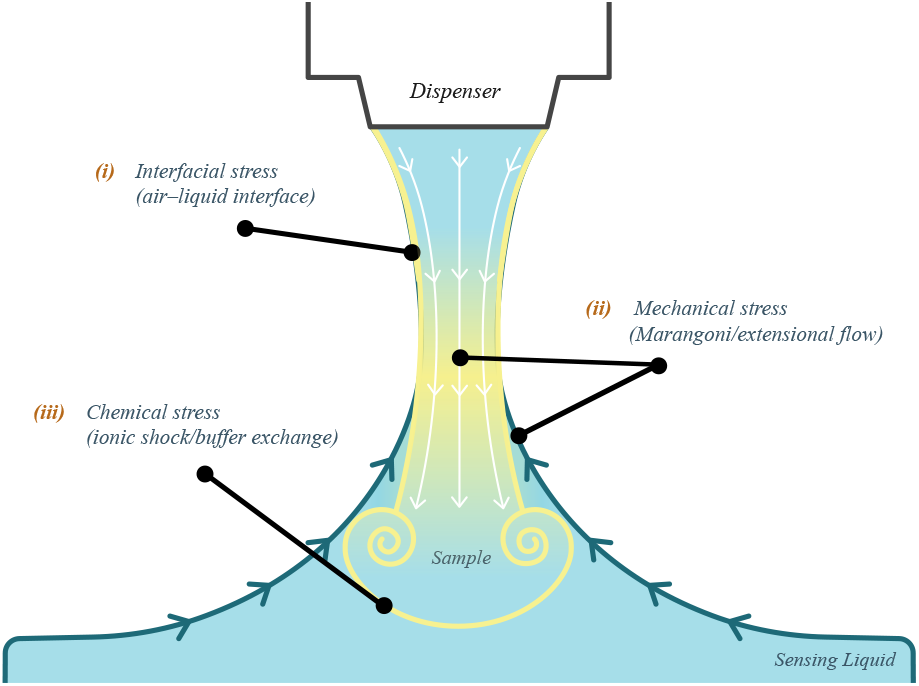
Time-ordered stresses imposed on the molecule during a single coalescence event. Three classes of stress act in defined temporal sequence. **(i) Interfacial stress** (∼4 s before contact): molecules adsorb to the expanding air–water interface, where adsorption imposes conformational stress as protein structure partially unfolds to maximize interfacial contact. **(ii) Mechanical stress** (*t* ≈ 0–20 ms): at rupture and during neck thinning, bulk extensional flow stretches soluble assemblies, the contracting interface compresses the adsorbed layer in-plane, and a surface-tension gradient drives Marangoni flow along the neck. **(iii) Chemical stress** (*t* ≈ 0–20 ms, concurrent with mechanical): surfactant and salt from the sensing-liquid reservoir mix into the molecule-bearing droplet at the contact boundary, shifting ionic strength, surfactant concentration, and chemical environment. The same classes of stress are encountered throughout a therapeutic’s life cycle (manufacturing, formulation, *in vivo*); the body of the Discussion details candidate mechanisms by which they may couple to produce the gel-like signature observed in Figure 6.

The gel-like signature we observe is consistent with several candidate contributions to the coalescence response: viscoelastic film formation during droplet growth [4, 5, 6, 28, 29, 30], extensional stretching of soluble assemblies [23, 24, 27, 31], and compression-driven unfolding of the adsorbed layer [3, 7, 32, 33]. The data here do not distinguish between these, and the time-ordered nature of the coalescence allows that more than one may contribute. Distinguishing them requires reconfiguring one stress at a time: as a concrete example, pre-loading the air– liquid interface with a competing surfactant added to the droplet itself should suppress the *σ*_*τ*_ response under the interfacial-film mechanism but not under either of the bulk alternatives. Such reconfigurations are the subject of ongoing work.

The instrument is configured so that complex behaviours of this kind appear in its recorded waveforms. The gel-like coalescence response we identify here is what we found by applying the instrument to antibody developability; we expect that other modalities (Apoha is extending the instrument to nanobodies and bispecifics) and other formulations will produce distinct emergent signatures, identifiable by the same top-down feature search that produced VIBE1.

Several limitations apply. The *R*^2^ = 0.92 between *σ*_*τ*_ and VIBE1 does not formally bound the residual ∼8%; whether it carries secondary information or nuisance variation needs larger replicate designs. The imaging set spans the dataset’s VIBE1 range but is too small (12 drops) to characterise the population-level distribution of *λ* across the 235 antibodies. We report *R*(*t*) and *H*(*t*) as kinematic observables of the assay geometry, with *λ* as its kinematic decay timescale rather than a constitutive relaxation time [34]. None of these affect the central result: *σ*_*τ*_ is a per-event readout of the antibody’s propensity to undergo a transient gel-like transition during coalescence, and that variability is what VIBE1 captures.

## Materials and Methods

### Sample preparation and measurement protocol

The 235-antibody dataset and full liquid-state-machine measurement protocol are described in Thomas *et al*. [1]; we summarise the aspects relevant here. All 235 are therapeutic monoclonal antibodies that have reached at least Phase 1 clinical trials, sourced from Sino Biological in PBS at pH 7.4 and diluted to 0.1 mg mL^−1^. Each sample was injected (80 µL) into a continuous PBS mobile-phase flow delivered by HPLC to the dispenser, deposited drop-by-drop (∼5 µL per drop, ∼100 drops per titration over ∼400 s) onto a proprietary near-thermodynamic-transition sensing liquid held near the critical micellisation concentration of an anionic surfactant. The instrument’s response channels were recorded ellipsometrically. Samples were tested in randomised batches of 20, in triplicate, at 22 ± 2°C, with sirukumab and trastuzumab as in-batch high- and low-VIBE1 controls. For concentration-response studies, sirukumab and trastuzumab were measured at five nominal concentrations (0.04, 0.06, 0.08, 0.10, 0.25 mg mL^−1^); BSA at 0.10, 0.20, 0.50 mg mL^−1^; PBS as the shared *c* = 0 baseline. All proteins were diluted from stock into the same PBS buffer. Trough and HPLC flow path were cleaned with water and PBS between titrations; the per-titration *τ*_*i*_ baseline measured over drops 0–9 (before plug arrival) returned to the buffer value at the start of each new measurement, indicating no carryover at the level of the bulk pinch-off time.

### Data extraction and statistical analysis

For each drop, the pinch-off time *τ*_*i*_ was identified from the upstream coalescence-sensor channel as the time of the secondary peak in the smoothed, standardised time-derivative of the signal: this peak corresponds to bridge rupture, while the first peak corresponds to the initial coalescence transient. Times are reported relative to the detector intensity minimum. The per-titration *σ*_*τ*_ is the standard deviation of *τ*_*i*_ across drops 20–40, the analysis window used by VIBE1. For analyses linking *σ*_*τ*_ to VIBE1, drops were excluded from the *σ*_*τ*_ calculation if their late *S*_Y_ pulse (generated at pinch-off and arriving with a ∼4 ms propagation delay) fell outside the 104–140 ms VIBE1 window: VIBE1 is computed only over that window, so out-of-window drops cannot contribute to a meaningful link. The exclusion removes 22 of ∼24,800 drops across the dataset (0.09%). For Figure 2D, *S*_Y_ traces from successive drops within a titration were time-shifted so that their *τ*_*i*_ values coincided; the residual ∼4 ms offset was estimated from cross-correlation of late-arriving pulses across drops. The dataset-level *R*^2^ between *σ*_*τ*_ and VIBE1 was obtained by linear regression on the 1182-titration dataset. Per-antibody Spearman *ρ* between each developability panel assay and either VIBE1 or per-antibody mean *σ*_*τ*_ was computed across antibodies with non-null readings. Hill curves overlaid on concentration-response data are descriptive envelopes, with 95% confidence bands from a 2,000-iteration parametric bootstrap; the data are insufficient to constrain a half-saturation concentration or Hill exponent quantitatively.

### High-speed imaging and image analysis

Coalescence events were imaged at 10,000 fps in transmitted-light bright-field, synchronised to droplet deposition, with the dispenser tip (1.59 mm OD; 26.8 µm/pixel calibration) in the upper field of view. Recording was triggered at drop 25 of an otherwise standard injection, capturing 2–3 consecutive drops per video. The analysis set comprises 12 drops: 1 water, 2 trastuzumab, 2 galiximab, and 7 fresolimumab. Each frame was bilinearly upsampled by a factor of three, Gaussian-smoothed, Otsu-thresholded into a binary droplet mask, and morphologically closed along the dispenser axis to bridge thin necks. The neck radius *R* is half the minimum mask width below the dispenser tip; the bottom edge of the upper droplet, post-rupture, is indexed in the same coordinate system as height *H*. The 3× upsample puts an effective floor of ∼5 µm (±0.2 native px) on *R*. The droplet–substrate contact frame was identified as the first frame after the recording trigger at which the smoothed mean image intensity dropped more than 5 standard deviations below the pre-contact baseline mean. The same threshold produces a consistent onset across the 12 imaged drops (Supplementary Fig. S4). The bridge-rupture frame was identified as the secondary peak in the smoothed time-derivative of the same mean-intensity trace; this matches the rupture event observed directly in the frames to within one video frame. The pre-rupture neck radius from *t* = *R*_max_ to *t* = *t*_rupture_ was fit by least squares to *R*(*u*) = *R*_∞_ + *R*_*e*_ (*e*^*u/*3*λ*^ − 1) with *u* = *t*_rupture_ − *t*, with bounds *R*_∞_ ∈ [0, 500] µm, *R*_*e*_ ∈ [0, 5000] µm, and *λ* ∈ [0.05, 50] ms. The form is a phenomenological exponential approach to a finite rupture radius; it is not the canonical Entov–Hinch / Anna– McKinley self-similar form, and *λ* is reported as a kinematic decay constant of this geometry, not as a polymer relaxation time [34]. Each drop’s post-rupture height trace *H*(*t*) is referenced to a per-drop equilibrium baseline *H*_eq_ estimated from late-time data. A drop is classified as oscillatory if the detrended trace executes a negative excursion of at least 8% of its positive excursion within 50 ms after rupture, and non-oscillatory otherwise. Per-drop diagnostic plots (frame thumbnails, *R*(*t*) and *H*(*t*), and the mean-intensity trace with its standardised time-derivative) are provided for every drop in the imaging set in Supplementary Fig. S3. The 12 underlying high-speed videos (10,000 fps, played back at reduced rate from −5 ms to +55 ms relative to contact) are provided as Supplementary Videos 1–12; the mapping from each video to its sample, replicate, and event-frame indices is given in Supplementary Table S1. Correlations involving high-speed imaging parameters are reported descriptively given the imaging sample size.

## Acknowledgements

We are grateful to Peter Tessier and Nimish Gera for helpful discussions and feedback during the course of this work.

## References

[1] Thomas AN, St John AN, Crames M, et al. (2026) Behavioural profiling of therapeutic antibodies via non-equilibrium interfacial wave dynamics. Manuscript submitted to bioRxiv (6 May 2026).

[2] Li J, et al. (2019) Interfacial stress in the development of biologics: fundamental understanding, current practice, and future perspective. AAPS J 21(3):44.

[3] Dobson J, Kumar A, Willis LF, et al. (2017) Inducing protein aggregation by extensional flow. Proc Natl Acad Sci USA 114(18):4673–4678.

[4] Kanthe A, et al. (2021) No ordinary proteins: adsorption and molecular orientation of monoclonal antibodies. Sci Adv 7(35):eabg2873.

[5] Wood CV, et al. (2023) Antibodies adsorbed to the air–water interface form soft glasses. Langmuir 39(22):7775–7782.

[6] Ruane S, et al. (2019) Interfacial adsorption of a monoclonal antibody and its Fab and Fc fragments at the oil/water interface. Langmuir 35(42):13543–13552.

[7] Willis LF, et al. (2018) Using extensional flow to reveal diverse aggregation landscapes for three IgG1 molecules. Biotechnol Bioeng 115(5):1216–1225.

[8] Grigolato F, Arosio P (2020) Synergistic effects of flow and interfaces on antibody aggregation. Biotechnol Bioeng 117(2):417–428.

[9] Pérez-Gil J (2008) Structure of pulmonary surfactant membranes and films: the role of proteins and lipid–protein interactions. Biochim Biophys Acta 1778(7-8):1676–1695.

[10] Schneider SW, Nuschele S, Wixforth A, Gorzelanny C, Alexander-Katz A, Netz RR, Schneider MF (2007) Shear-induced unfolding triggers adhesion of von Willebrand factor fibers. Proc Natl Acad Sci USA 104(19):7899–7903.

[11] Birtalan S, Zhang Y, Fellouse FA, Shao L, Schaefer G, Sidhu SS (2008) The intrinsic contributions of tyrosine, serine, glycine and arginine to the affinity and specificity antibodies of. J Mol Biol 377(5):1518–1528.

[12] Waibl F, et al. (2021) Conformational ensembles of antibodies determine their hydrophobicity. Biophys J 120(1):143–157.

[13] Jain T, et al. (2017) Biophysical properties of the clinical-stage antibody landscape. Proc Natl Acad Sci USA 114(5):944–949.

[14] Jain T, Boland T, Vásquez M (2023) Identifying developability risks for clinical progression of antibodies using high-throughput in vitro and in silico approaches. mAbs 15(1):2200540.

[15] Mieczkowski C, Zhang X, Lee D, Nguyen K, Lv W, Wang Y, Zhang Y, Way J, Gries JM (2023) Blueprint for antibody biologics developability. mAbs 15(1):2185924.

[16] Shieh IC, Patel AR (2015) Predicting the agitation-induced aggregation of monoclonal antibodies using surface tensiometry. Mol Pharm 12(9):3184–3193.

[17] Eggers J (1997) Nonlinear dynamics and breakup of free-surface flows. Rev Mod Phys 69(3):865–929.

[18] Eggers J, Sprittles JE, Snoeijer JH (2025) Coalescence dynamics. Annu Rev Fluid Mech 57:61–87.

[19] Tirtaatmadja V, McKinley GH, Cooper-White JJ (2006) Drop formation and breakup of low viscosity elastic fluids: effects of molecular weight and concentration. Phys Fluids 18(4):043101.

[20] Pawar AB, Caggioni M, Ergun R, Hartel RW, Spicer PT (2011) Arrested coalescence in Pickering emulsions. Soft Matter 7(17):7710–7716.

[21] Pawar AB, Caggioni M, Hartel RW, Spicer PT (2012) Arrested coalescence of viscoelastic droplets with internal microstructure. Faraday Discuss 158:341–350.

[22] Sharma V, Jaishankar A, Wang YC, McKinley GH (2011) Rheology of globular proteins: apparent yield stress, high shear rate viscosity and interfacial viscoelasticity of bovine serum albumin solutions. Soft Matter 7(11):5150–5160.

[23] Anna SL, McKinley GH (2001) Elasto-capillary thinning and breakup of model elastic liquids. J Rheol 45(1):115–138.

[24] Entov VM, Hinch EJ (1997) Effect of a spectrum of relaxation times on the capillary thinning of a filament of elastic liquid. J Non-Newtonian Fluid Mech 72(1):31–53.

[25] Lin GL, Pathak JA, Kim DH, Carlson M, Riguero V, Kim YJ, Buff JS, Fuller GG (2016) Interfacial dilatational deformation accelerates particle formation in monoclonal antibody solutions. Soft Matter 12(14):3293–3302.

[26] Scriven LE, Sternling CV (1960) The Marangoni effects. Nature 187(4733):186–188.

[27] Dekker PJ, Hack MA, Tewes W, Datt C, Bouillant A, Snoeijer JH (2022) When elasticity affects drop coalescence. Phys Rev Lett 128(2):028004.

[28] Koepf E, Eisele S, Schroeder R, Brezesinski G, Friess W (2018) Notorious but not understood: how liquid–air interfacial stress triggers protein aggregation. Int J Pharm 537(1– 2):202–212.

[29] Mehta SB, Lewus LM, Bee JS, et al. (2015) Gelation of a monoclonal antibody at the silicone oil–water interface and subsequent rupture of the interfacial gel results in aggregation and particle formation. J Pharm Sci 104(4):1282–1290.

[30] Bazazi P, Stone HA, Hejazi SH (2023) Dynamics of droplet pinch-off at emulsified oil–water interfaces: interplay between interfacial viscoelasticity and capillary forces. Phys Rev Lett 130(3):034001.

[31] McKinley GH (2005) Visco-elasto-capillary thinning and break-up of complex fluids. In: Rheology Reviews 2005, British Society of Rheology, Aberystwyth, pp. 1–49.

[32] Hughes MDG, Hanson BS, Cussons S, Mahmoudi N, Brockwell DJ, Dougan L (2021) Control of nanoscale in situ protein unfolding defines network architecture and mechanics of protein hydrogels. ACS Nano 15(7):11296–11308.

[33] Hughes MDG, Cussons S, Mahmoudi N, Brockwell DJ, Dougan L (2022) Tuning protein hydrogel mechanics through modulation of nanoscale unfolding and entanglement in post-gelation relaxation. ACS Nano 16(7):10667–10678.

[34] Hu N, Hwang J, Ruangkriengsin T, Stone HA (2025) Revealing actual viscoelastic relaxation times in capillary breakup. Phys Rev Lett 135(4):048201. doi:10.1103/2jz7-4w4k

