## Supplementary Information for "Millisecond nonlinear state changes during droplet coalescence identify therapeutic-antibody developability liabilities"

This Supplementary Information accompanies the manuscript.

#### Supplementary Videos

Supplementary Videos 1–12 are the raw high-speed-video recordings underlying the per-drop kinematic analysis (Methods; main-text Figures 4–6; Supplementary Figs. S1–S4). All clips were acquired at 10,000 fps in transmitted-light bright-field on the same instrument as the dataset measurements, and are presented at reduced playback rate. The contact frame and bridge-rupture frame for each clip are listed below; both refer to the original 10,000 fps acquisition.

| Video | Sample | Drop no. | Contact frame | Rupture frame |
|---|---|---|---|---|
| 1 | water | 0 | 961 | 1083 |
| 2 | trastuzumab | 0 | 966 | 1095 |
| 3 | trastuzumab | 1 | 969 | 1097 |
| 4 | galiximab | 0 | 961 | 1094 |
| 5 | galiximab | 1 | 963 | 1113 |
| 6 | fresolimumab | 0 | 967 | 1224 |
| 7 | fresolimumab | 1 | 970 | 1137 |
| 8 | fresolimumab | 2 | 968 | 1192 |
| 9 | fresolimumab | 3 | 965 | 1139 |
| 10 | fresolimumab | 4 | 969 | 1223 |
| 11 | fresolimumab | 5 | 968 | 1124 |
| 12 | fresolimumab | 6 | 968 | 1178 |

**Supplementary Table S1: Index of supplementary videos.** Each row corresponds to one of the 12 imaged coalescence events. `Drop no.` matches the per-drop identifier used in Supplementary Fig. S3. Contact and rupture frames are indices into the original 10,000 fps acquisition.

#### Per-sample high-speed-video traces

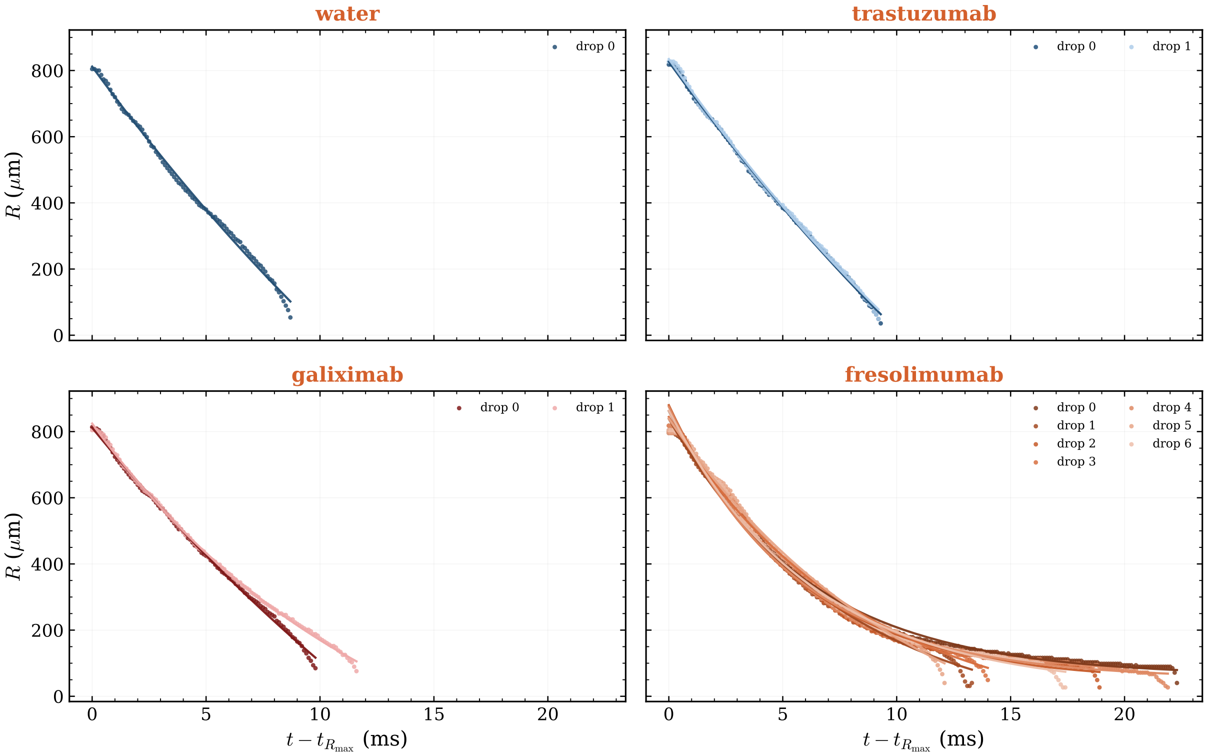


**Supplementary Fig. S1: Pre-rupture neck radius $R(t)$ for every imaged drop, one panel per sample.** Each panel shows all imaged drops of the labelled sample, anchored at $t = R_{\max}$. Solid points: smoothed-mask data; solid lines: per-drop elasto-capillary fit $R(u) = R_\infty + R_e(e^{u/3\lambda} - 1)$ with $u = t_{\text{rupture}} - t$ (Methods). The four panels share axes. The early portion of every $R(t)$ curve, water and antibodies alike, overlaps within imaging precision; divergence appears only as the neck narrows, with the largest deceleration in the long-pinch fresolimumab drops.

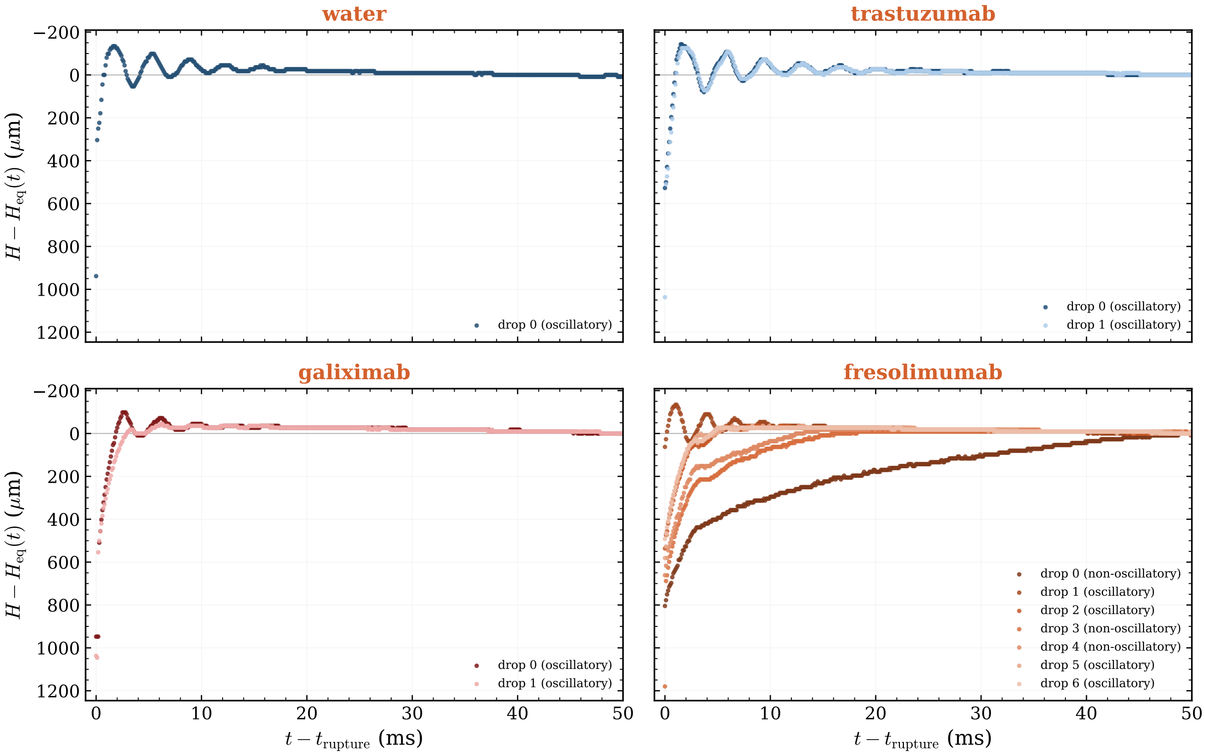


**Supplementary Fig. S2: Post-rupture detrended height $H - H_{eq}(t)$ for every imaged drop, one panel per sample.** Each panel shows all imaged drops of the labelled sample. Each drop's recoil class (oscillatory vs. non-oscillatory) is given in the legend. The reference equilibrium $H_{eq}$ is the per-drop median $H_{raw}$ in a $\pm 2$ ms window centred on $t_{rupture} + 50$ ms, so each detrended trace passes through zero at the anchor by construction (Methods). The $y$-axis is inverted so that retraction (upper drop sitting higher than equilibrium) appears as positive deflection. Water, trastuzumab, and galiximab drops oscillate freely about the equilibrium baseline and settle to it; some of the long-pinch fresolimumab drops persist at a non-spherical shape through the imaging window.

### Annotated per-drop high-speed-video frames

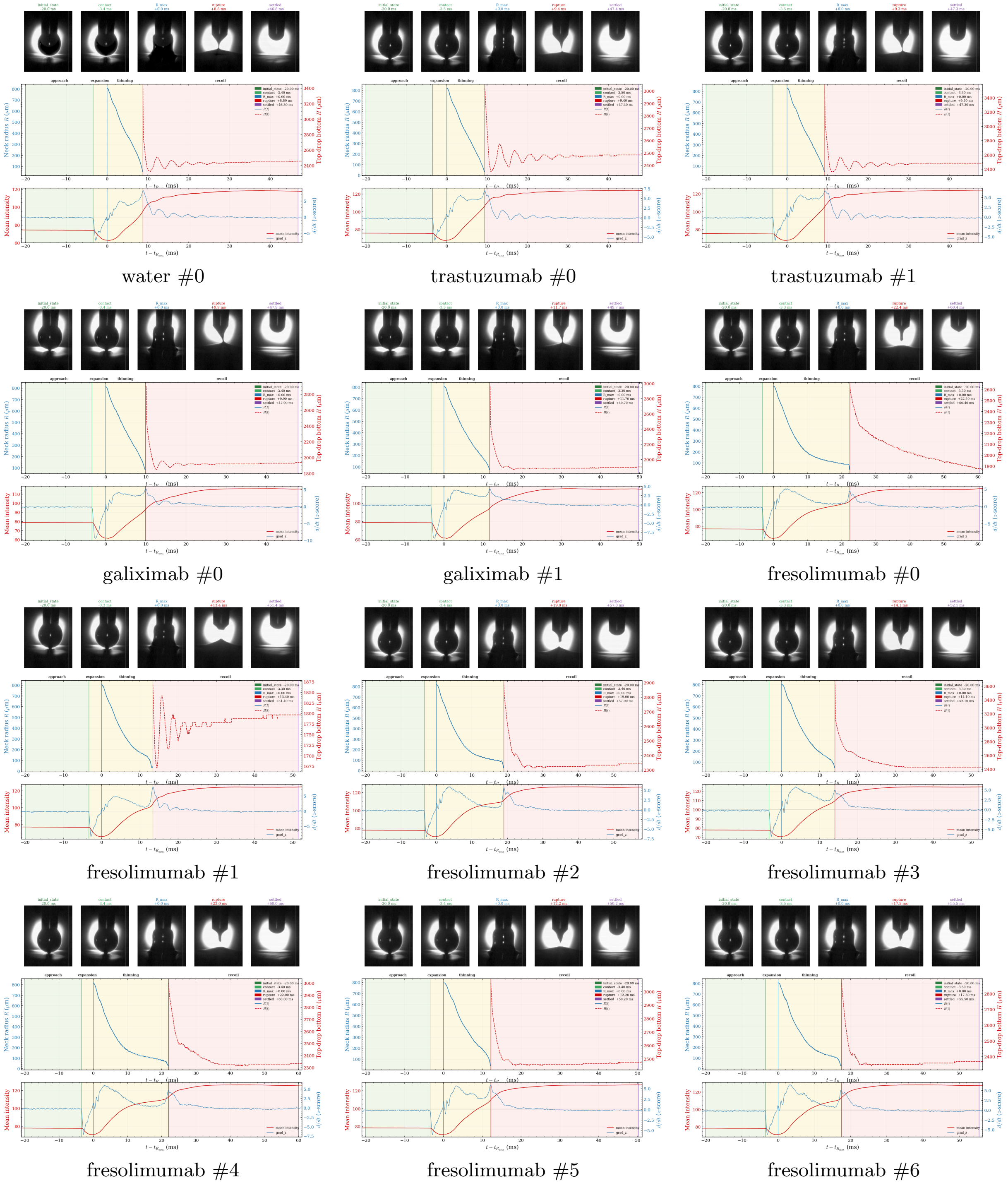


**Supplementary Fig. S3: Per-drop diagnostic plots for every drop in the high-speed-imaging set.** Each panel shows, top to bottom: five frame thumbnails labelled by deposition stage (initial state, contact, $R_{\max}$, rupture, settled); the pre-rupture neck radius $R(t)$ (blue) and post-rupture upper-drop bottom edge $H(t)$ (red, dashed) on a twin-axis with phase shading; and the frame-by-frame mean image intensity (red) with its standardised time-derivative (blue), used for event-frame identification (Methods).

### Onset-alignment diagnostic for the high-speed-imaging set

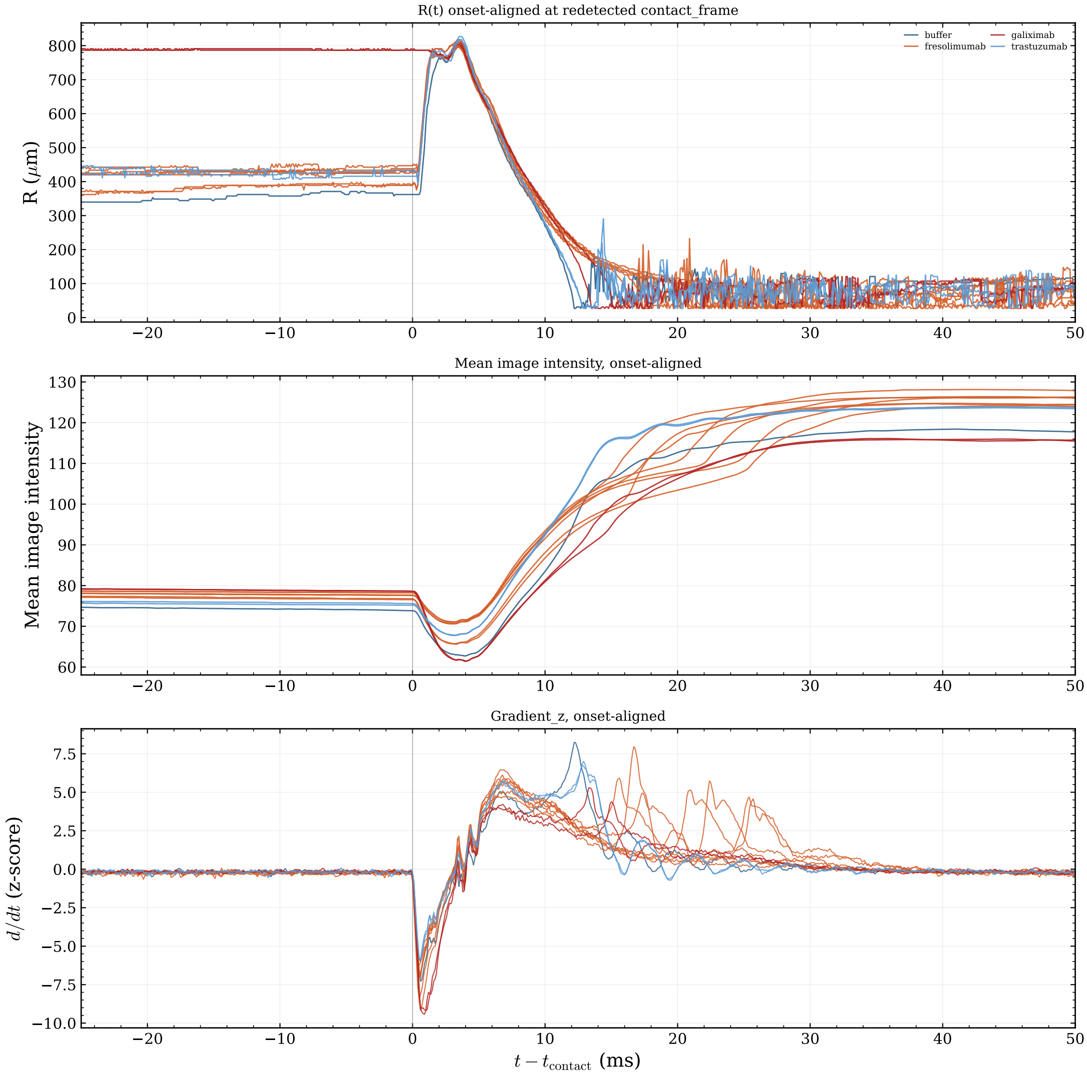


**Supplementary Fig. S4: The 5$\sigma$ contact-frame detection produces a consistent onset across the high-speed-imaging set.** All 12 imaged drops, anchored at the redetected contact frame ($t = 0$). Top: pre-rupture neck radius $R(t)$. Middle: frame-by-frame mean image intensity. Bottom: standardised, smoothed time-derivative of the mean intensity. The contact frame is identified as the first frame at which the smoothed mean intensity drops more than 5 standard deviations below its pre-contact baseline; the bridge-rupture frame is identified as the secondary peak in the time-derivative (Methods). Every drop's intensity drops in the same way at $t = 0$ and the secondary derivative peak appears in a tight window across drops, validating the event-frame identification used for the per-drop $R(t)$, $H(t)$ extractions.

#### Per-antibody robustness of $\sigma_\tau$–VIBE1

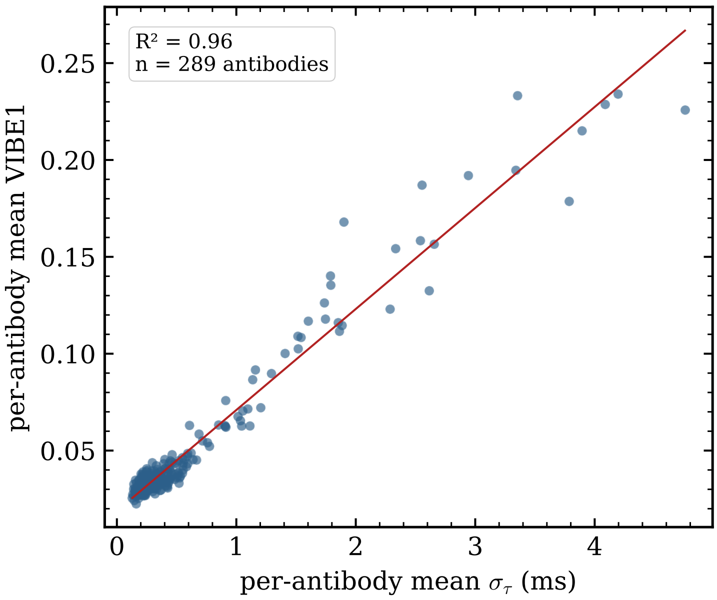


**Supplementary Fig. S5: Per-antibody mean $\sigma_\tau$ vs. per-antibody mean VIBE1.** The cohort scatter of main-text Figure 3A is at the per-titration level; here both quantities are averaged across replicate titrations of each antibody. Linear regression on antibodies with non-null readings gives Pearson $R^2 = 0.96$ ($n = 289$). The relationship between $\sigma_\tau$ and VIBE1 is therefore preserved when within-antibody replicate variability is averaged out.
